# Towards Resolving and Redefining Amphipyrinae (Lepidoptera, Noctuoidea, Noctuidae): a Massively Polyphyletic Taxon

**DOI:** 10.1101/271478

**Authors:** Kevin L. Keegan, J. Donald Lafontaine, Niklas Wahlberg, David L. Wagner

## Abstract

Amphipyrinae have long been a catchall taxon for Noctuidae, with most members lacking discernible morphological synapomorphies that would allow their assignment to one of the many readily diagnosable noctuid subfamilies. Here data from seven gene regions (>5,500 base pairs) for more than 120 noctuid genera are used to infer a phylogeny for Amphipyrinae and related subfamilies. Sequence data for 57 amphipyrine genera—most represented by the type species of the genus—are examined. Presented here are: the first large-scale molecular phylogenetic study of Amphipyrinae and largest molecular phylogeny of Noctuidae to date; several proposed nomenclatural changes for well supported results; and the identification of areas of noctuid phylogeny where greater taxon sampling and/or genomic-scale data are needed. Adult and larval morphology, along with life history traits, for taxonomic groupings most relevant to the results are discussed. Amphipyrinae are significantly redefined; many former amphipyrines, excluded as a result of these analyses, are reassigned to other noctuid subfamily-level taxa. Four genera, *Chamaeclea* Grote, *Heminocloa* Barnes & Benjamin, *Hemioslaria* Barnes & Benjamin, and *Thurberiphaga* Dyar are transferred to the tribe Chamaecleini Keegan & Wagner **New Tribe** in Acontiinae. Stiriina is elevated to Stiriinae **Revised Status**, Grotellina is elevated to Grotellinae **Revised Status**, and Annaphilina is elevated to Annaphilini **Revised Status**. *Acopa* Harvey is transferred to Bryophilinae, *Aleptina* Dyar is transferred to Condicinae, *Leucocnemis* Hampson and *Oxycnemis gracillinea* (Grote) are transferred to Oncocnemidinae, *Nacopa* Barnes & Benjamin is transferred to Noctuinae, and *Narthecophora* Smith is transferred to Stiriinae. *Azenia* Grote (and its subtribe Azeniina), *Cropia* Walker, *Metaponpneumata* Möschler, *Sexserrata* Barnes & Benjamin, and *Tristyla* Smith are transferred to Noctuidae *incertae sedis*. *Hemigrotella* Barnes & McDunnough (formerly in subtribe Grotellina) is retained in Amphipyrinae.

This published work has been registered in ZooBank, http://zoobank.org/urn:lsid:zoobank.org:pub:4A140782-31BA-445A-B7BA-6EAB98ED43FA

## INTRODUCTION

Amphipyrinae have long been a taxon of uncertain identity. In the case of some its tribes and subtribes, placement within the subfamily has been simply a matter of nomenclatural convenience (Poole, 1995). In essence, Amphipyrinae became a “junk drawer” for Noctuidae: a repository for taxa lacking the characters other subfamilies (Poole, 1995; Kitching, 1984; Kitching & Rawlins, 1998; Fibiger & Lafontaine, 2005). As a consequence, taxonomic concepts of what is and is not an amphipyrine have varied greatly through time, across continents, and among workers.

Hampson’s (1898-1913) world classification of noctuids provided an expansive concept of Amphipyrinae, rendering it a massive group of morphologically heterogeneous moths accounting for nearly half of the world’s described noctuid genera (sensu Lafontaine & Schmidt, 2010) at the time (Kitching, 1984). When Poole (1989) published his catalog of the world’s noctuid genera, several groups had been removed from Amphipyrinae (e.g. Acronictinae), but his Amphipyrinae still included over 500 genera. Kitching & Rawlins (1998) were so vexed by what is and what is not an amphipyrine that they restricted membership to just the nominate genus, *Amphipyra* Ochsenheimer.

In North America, many noctuid collections, Internet resources, and taxonomic literature are organised according to Franclemont & Todd’s (1983) checklist of Nearctic moths found north of Mexico. Their concept of Amphipyrinae included more than five dozen genera presently classified as Noctuinae; many genera now assigned to Balsinae, Bryophilinae, Condicinae, Eriopinae, Metoponiinae; more than two dozen “unassociated genera,” most of which were reclassified by Lafontaine & Schmidt (2010, 2015) into other subfamilies; as well as a few erebids and a nolid! In Africa, Australia, Japan, and other parts of Asia, the subfamily’s limits remain more Hampsonian and nebulous, overlapping with Acronictinae, Noctuinae, and other subfamilies (Hampson, 1898-1913; Edwards, 1996).

Subsequent morphological and molecular studies challenged the classifications of Franclemont & Todd (1983) and Kitching & Rawlins (1998), dramatically reshuffling the contents of Amphipyrinae and other noctuid subfamilies. Fibiger & Lafontaine’s (2005) reclassification of Noctuoidea relied on morphological characters to redefine families and subfamilies using known character systems such as the position of the clasper in the male genitalia and features of the tympanum, as well as novel character systems such as the presence of setae on the scaphium and whether the lateral stripe of larvae continued around the anal plate or dropped down the anal proleg. In their treatment, Amphipyrinae were restricted to just the genus *Amphipyra* plus *Phidrimana* Kononenko and *Pyrois* Hübner. Based on their assessment, Amphipyrinae, Psaphidinae, and Stiriinae exhibited mixtures of primitive and derived states and accordingly were grouped near each other in the middle of their phylogenetic sequence of subfamilies. Wagner *et al.* (2008) recommended subsuming Psaphidinae into Amphipyrinae, as a tribe, based on shared larval characters (e.g., head retracted into prothorax and A8 being sharply angulate) and male genitalic features (e.g., finger-like ampulla and vesica with numerous spike-like cornuti). Lafontaine and Schmidt’s (2010) concept of Amphipyrinae removed more than 150 of Franclemont & Todd’s (1983) amphipyrine genera, and included Psaphidini and Stiriini. The latter tribe Poole (1995), Kitching & Rawlins (1998), Mitchell *et al.* (2006), and others had previously supported as belonging in a separate subfamily.

Recent molecular phylogenetic studies also added to the sea-change of subfamilial taxonomic classification within Noctuidae. Mitchell *et al.* (2006) sampled broadly across noctuid subfamilies (sensu Lafontaine & Schmidt, 2010) including approximately 100 noctuid genera with special emphasis on subfamilies originating from shallower nodes in their noctuid phylogeny (e.g. Heliothinae and Noctuinae). Studies by Zahiri *et al.* (2011, 2012, 2013) focused on family relationships within Noctuoidea, as well as clarifying relationships among several noctuid subfamilies originating from deeper nodes. Rota *et al.* (2016) examined noctuid subfamilial relationships in and around Acronictinae, a subfamily thought to be closely related to Amphipyrinae. Regier *et al.* (2017) assessed subfamilial relationships across Noctuidae, corroborating previous studies on subfamilial relationships and finding strong support for many deep nodes within Noctuidae. Although these studies clarified many subfamilial relationships across Noctuidae, no previous study has sampled extensively in Amphipyrinae—one of the remaining great unknowns of noctuid classification.

The guide for taxon sampling in this study was the North American (north of Mexico) Noctuoidea checklist of Lafontaine & Schmidt (2010, 2015). Their concept of the Amphipyrinae consisted of approximately 225 species in 73 genera parsed out among three tribes, eight subtribes, and an *incertae sedis* group, with the majority of this diversity occurring in deserts and other aridlands of southwestern North America. In terms of generic diversity, these 73 genera represent approximately 75% of the world’s amphipyrine generic diversity (JDL unpublished data). By comparison, Amphipyrinae in Europe include only nine genera, with three of these genera shared with the North American fauna (Fibiger & Hacker, 2004)

This preliminary study of the Amphipyrinae uses 5,508 base pairs from mitochondrial and nuclear genes to test the monophyly of predominantly Nearctic amphipyrines. As much as possible, type species of genera were included. Although several amphipyrine genera were not included in this study, it represents the most comprehensive phylogenetic assessment of the subfamily and the Noctuidae to date with more than 120 noctuid genera sampled, representing 21 recognised subfamilies. In this effort, nomenclatural recommendations are limited to well supported results, and areas of noctuid phylogeny, proximate to the Amphipyrinae, are identified where greater taxon sampling is needed. Much discussion is given to providing adult and larval characters associated with the major clades whose content is affected by the results of this study.

## METHODS

### Taxon sampling

Sequence data for 63 species representing 61 noctuid genera were generated, few of which had been included in previous molecular phylogenetic studies. Fifty-seven of the 76 Nearctic genera in Amphipyrinae, as circumscribed by Lafontaine & Schmidt (2010, 2015), were included; representing all three amphipyrine tribes, all eight subtribes, and all seven *incertae sedis* genera (see Table S1 in supplementary materials). Forty-seven of the 57 amphipyrine genera were represented by their type species. For amphipyrine genera for which the type species was not sampled, morphologically similar and/or COI-proximate congeners were selected. Single specimens of each species were used. Collection and deposition information for voucher specimens newly collected for this study can be found in Table S1.

Data newly generated for this study were combined with the dataset published by Zahiri *et al.* (2013) as well as selected taxa representing independent lineages from Rota *et al.* (2016) (Table S1). These datasets represent all of the major lineages of Noctuidae sequenced to date, using the same genes as in this study (see Gene Sampling below), and serve as outgroups. Additional outgroups included members of the other noctuoid families and, in the case of Notodontidae, were used to root the tree.

### Gene Sampling

Seven genes were sampled, which in previous studies have been shown to be capable of resolving phylogenetic relationships of Lepidoptera at differing evolutionary depths: COI, EF-1α, GAPDH, IDH, MDH, RpS5, and wingless (Cho *et al.*, 1995; Fang *et al.*, 1997; Mitchell *et al.*, 2006; Wahlberg & Wheat, 2008; Zahiri *et al.*, 2011, 2013; Rota *et al.*, 2016; Regier *et al.*, 2017). Both COI and EF-1α were sequenced in two parts making for a total of nine loci. CAD, which has been used to study the molecular systematics of noctuids in conjunction with the seven genes mentioned above (Zahiri *et al.*, 2011, 2013; Rota *et al.*, 2016), was abandoned due to its low amplification success during initial PCR runs.

### DNA Extraction, PCR Sequencing, and Alignment

All DNA extractions were done using the NucleoSpin Tissue 250 kit manufactured by Macherey-Nagel using 1-2 legs from each specimen. Once extracted, DNA was stored in a refrigerator at ~4° C until needed for PCR. The PCR profiles and primers outlined in Wahlberg & Wheat (2008) were used. PCR products were sent to Macrogen Europe Inc. (Amsterdam, the Netherlands) or Macrogen USA Inc. (Rockville, Maryland) for Sanger sequencing. For the majority of loci, single forward reads were used, although some fragmented PCR products required reverse reads. Sequence chromatograms were visually inspected for base call errors and heterozygous loci in Geneious^®^ 8.1.9 (http://www.geneious.com, Kearse *et al.*, 2012). Consensus sequences for dual-read loci were also generated in Geneious. To ensure sequences were attributed to the correct species, a local BLAST (Altschul *et al.*, 1990) search was conducted in Geneious to compare the manually named sequence files with the unnamed sequences from Macrogen. Sequences were then checked against sequences available in GenBank (NCBI Resource Coordinators, 2017) and BOLD (Ratnasingham & Hebert, 2007) to detect misdeterminations and contamination. After being exported to FASTA files, sequences were visually aligned to reference lepidopteran sequences for each locus using AliView version 1.18 (Larsson, 2014), and then concatenated using AMAS version 0.95 (Borowiec, 2016). Phylogenetic hypotheses were inferred for each locus to detect possible contamination. GenBank accession numbers for sequences can be found in Table S1.

### Phylogenetic Inference and Tree Visualization

The 567 newly generated sequences were analysed in conjunction with 810 published noctuoid sequences from Zahiri *et al.* (2011, 2013) and Rota *et al.* (2016). The concatenated alignment was partitioned by gene and by codon position, giving a total of 21 partitions. Phylogenetic hypotheses were inferred with RAxML using the RAxML BlackBox web-server (Stamatakis *et al.*, 2008), IQ-TREE (Nguyen *et al.*, 2015), and MrBayes v. 3.2.6 (Ronquist *et al.*, 2012) all using the CIPRES web server (Miller *et al.*, 2010).

For the RAxML analysis, in addition to searching for the maximum likelihood tree, a bootstrap (BS) analysis with 1000 replicates was performed. For the IQ-TREE analysis, a model finding (Kalyaanamoorthy *et al.*, 2017) as well as a partition finding (Chernomor *et al.*, 2016) procedure (command TESTNEWMERGE) were run prior to searching for the maximum likelihood tree. Clade support in the IQ-TREE analysis was assessed with 1000 replicates of ultrafast bootstrap (UF) (Hoang *et al.*, 2018) and 1000 replicates of SH-aLRT (SH) (Guindon *et al.*, 2010). For the MrBayes analysis, two independent runs of 10,000,000 generations were run, each with one cold and seven heated chains. Clade support was assessed with posterior probabilities (PP). For this study, results are considered well supported or with good support for RAxML when BS >= 70 (Hillis & Bull, 1993), IQ-TREE when UF>= 95 and SH >= 80 (Trifinopoulos & Minh, 2018), and MrBayes when PP >= 0.95.

The stationarity of MCMC parameters estimated with MrBayes was assessed with Tracer v 1.6.0 (Rambaut *et al.*, 2014). Tree files and alignments are available from the Dryad Digital Repository: https://doi.org/10.5061/dryad.qm2kg13. The R (R Core Team, 2017) package ggtree v1.10.5 (Yu *et al.*, 2017) in R Studio v 1.0.383 (R Studio Team, 2015) was used to visualise and annotate the trees. Further annotation was done using GIMP and Adobe Photoshop image-editing software.

### Morphological and Life History Assessment

Clade membership and topological positions of all amphipyrine genera were evaluated in terms of their male genital characters by JDL. At least one dissection was examined or newly prepared for most genera, and in all instances where a genus fell outside of Amphipyrinae, Metoponiinae, and Stiriinae as depicted in Fig. 2B. Likewise, phylogenetic positions were evaluated in terms of larval biology and morphology by DLW. Findings that reinforce or refute the molecularly inferred phylogenetic relationships are reported in the Discussion.

## RESULTS

The dataset consisted of concatenated sequences of 154 noctuoid species with a maximum of 5,508 sites for the combined seven gene regions (and nine loci)—2,009 (36.5%) of the sites were parsimony informative. On average, each taxon’s sequence data consisted of 25.1% missing or ambiguous sites. See Table S1 for sequence coverage by gene and taxon. No major signs of sequence contamination, no major conflicts among the three phylogenetic analyses, and no convergence problems in the Bayesian analysis were found. Although there was good support for many of the shallow nodes in the analysis, many deeper nodes underpinning inter-subfamilial relationships were not as well supported (a matter returned to in the Discussion). The topology of the RAxML analysis is presented in tree figures with nodal support indicated for bootstrap values greater than or equal to 70; nodal support values from the IQ-TREE and MrBayes analyses are included in relevant sections of the text.

Amphipyrinae proved to be surprisingly polyphyletic, with their genera supported as members of circa ten subfamily-level noctuid lineages (Fig. 1). A much restricted Amphipyrinae (Amphipyrinae s.s.) were suggested with over half of their species-level diversity belonging elsewhere in the Noctuidae (Figs 2A,B). Amphipyrinae s.s. consist largely of Lafontaine & Schmidt’s (2010) tribes Amphipyrini and Psaphidini (Fig. 3) along with the East Asian genus *Nacna* Fletcher. The clade was not well supported in the RAxML analysis (BS=62), but was well supported in the IQ-TREE (UF=99, SH=99.4) and MrBayes analyses (PP=0.972).

**Fig. 1.**
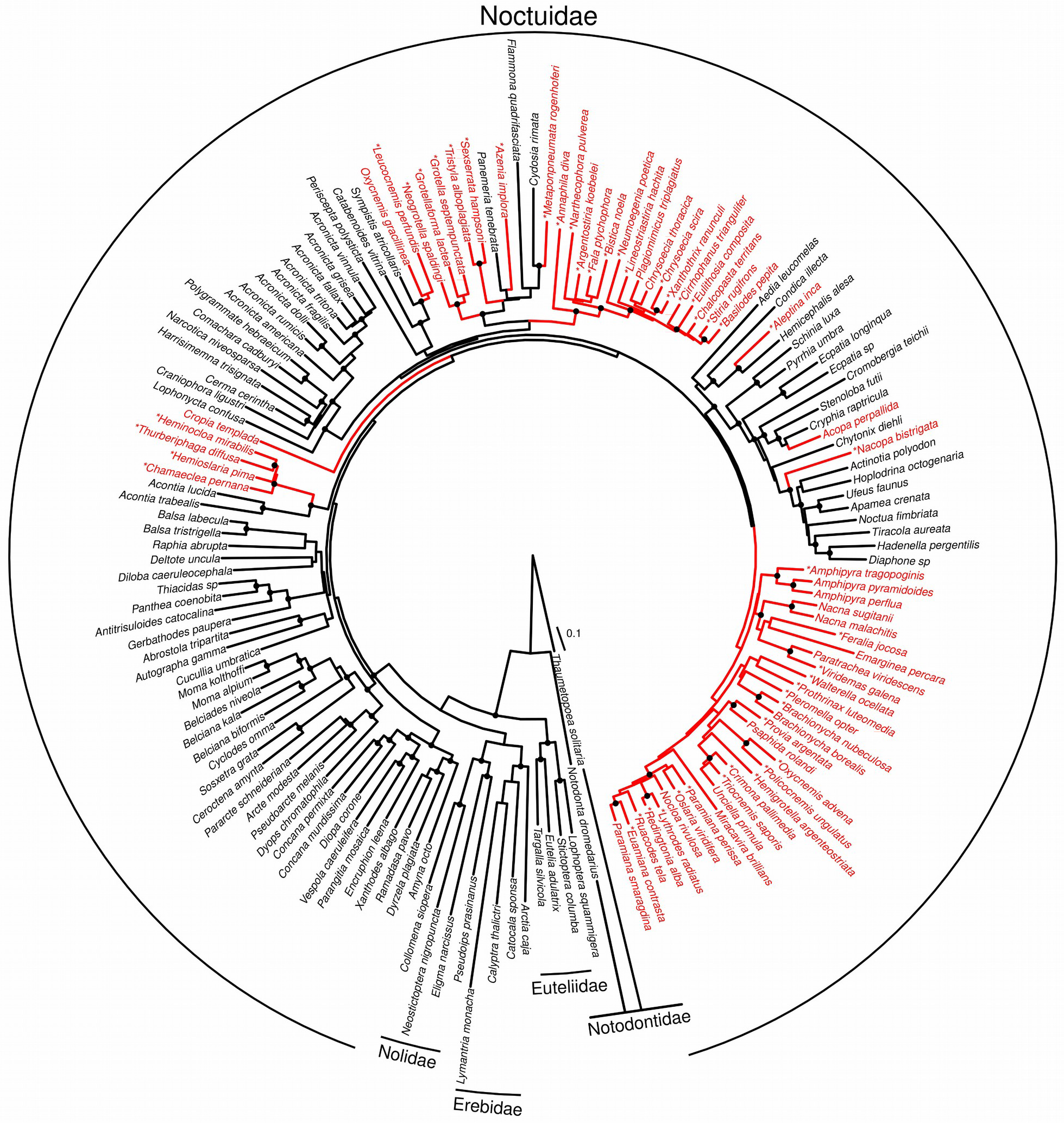
Results of seven-gene RAxML analysis. All Noctuoidea lineages in the dataset are shown. Lineages colored in red are classified as Amphipyrinae according to Lafontaine and Schmidt (2010, 2015). Scale bar shows expected substitutions per site. Nodes with bootstrap >= 70 are shown as black dots (see Methods for bootstrap details). Type species for amphipyrine genera are denoted with an asterisk.

**Figs 2A,B.**
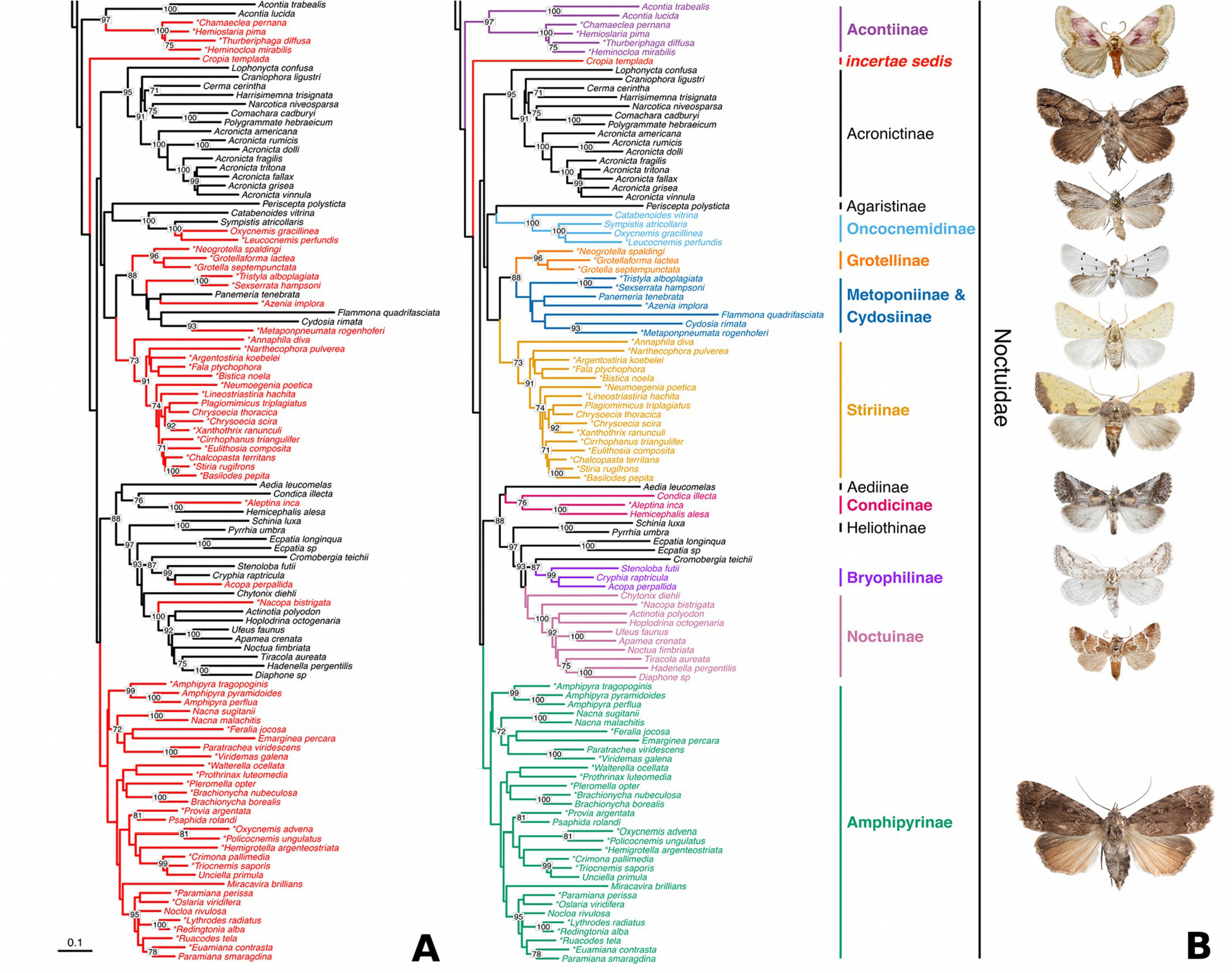
A, Results of seven-gene RAxML analysis: only the extent of the tree containing Amphipyrinae (sensu Lafontaine and Schmidt, 2010, 2015) lineages shown. Lineages colored in red are classified as Amphipyrinae. Bootstrap values >= 70 are displayed (see Methods for bootstrap details). Type species for amphipyrine genera denoted with an asterisk. B, Same analysis as in Fig 2A but with subfamily-level taxa that contain amphipyrine taxa (sensu Lafontaine and Schmidt, 2010, 2015) colored and labeled in bold. For each of the subfamily-level taxa that contain amphipyrines, a representative species used in the analysis is pictured near its position in the tree. From top to bottom: *Chamaeclea pernana* (Grote), *Cropia connecta* (Smith), *Oxycnemis gracillinea* (Grote), *Grotella septempunctata* Harvey, *Azenia implora* Grote, *Stiria rugifrons* Grote, *Aleptina inca* Dyar, *Acopa perpallida* Grote, *Nacopa bistrigata* (Barnes & McDunnough), *Amphipyrapyramidoides* Guenee. Images are (roughly) scaled to life size.

**Fig. 3.**
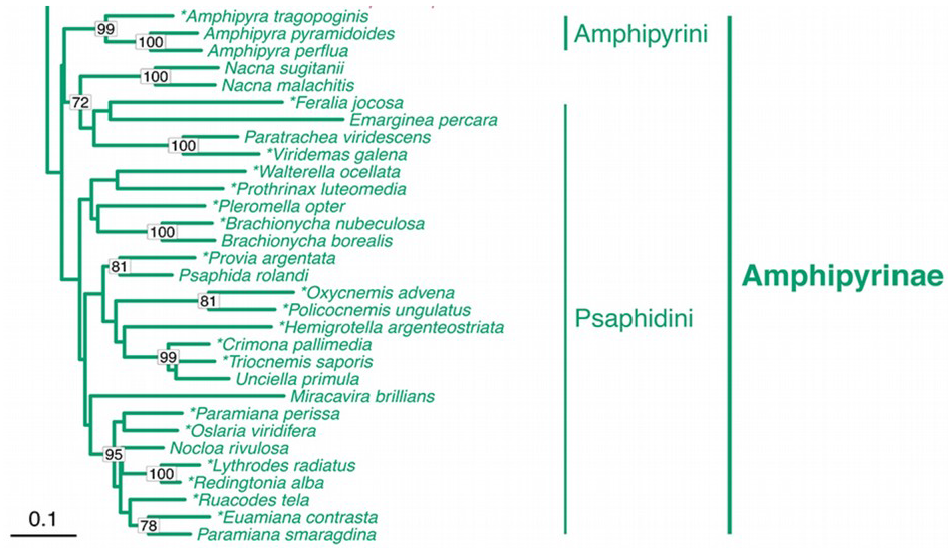
Subset of Fig 2B showing just Amphipyrinae s.s. with tribes (Amphipyrini and *Psaphidini) labeled. Nacna* Fletcher is not treated by Lafontaine and Schmidt (2010, 2015).

The amphipyrine tribe Stiriini was shown to be polyphyletic with much of its diversity spread across three subfamilies: Stiriinae **Revised Status** (BS=73, UF=99, SH=94.3, PP=0.998), Metoponiinae, and Grotellinae **Revised Status** (BS=96, UF=100, SH=100, PP=1.00) (Fig. 4). Grotellinae contain the genera of the former Grotellina, except *Hemigrotella* Barnes & McDunnough, which grouped within Amphipyrinae s.s. Stiriinae comprise two tribes: Stiriini **Revised Status** (BS=91, UF=100, SH=94.7, PP=0.912) and Annaphilini **Revised Status**. Stiriini contain, in large part, the contents of the former Stiriina, as well as *Narthecophora* Smith (formerly a member of the amphipyrine subtribe Azeniina) and two genera listed as *incertae sedis* in Stiriini by Lafontaine & Schmidt (2010): *Argentostiria* Poole and *Bistica* Dyar. Annaphilini contain *Annaphila* Grote and *Axenus* Grote (not included in this analysis). Stiriinae grouped sister to the clade containing Metoponiinae, Cydosiinae, and Grotellinae (BS=65, UF=99, SH=100, PP=0.979).

**Fig. 4.**
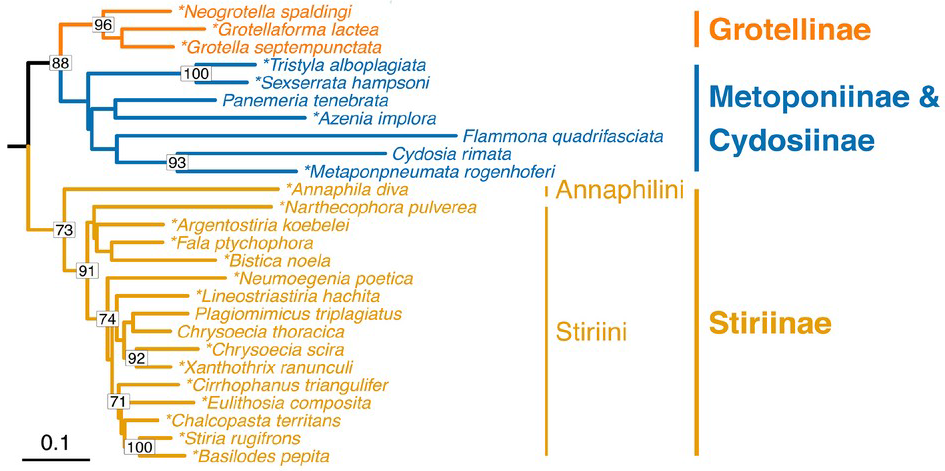
Subset of Fig 2B showing Grotellinae, Metoponiinae, Cydosiinae, and Stiriinae.

*Azenia* Grote (type genus of Azeniina) grouped within the clade containing Metoponiinae and Cydosiinae (BS=64, UF=99, SH=97.1, PP=0.959). Also clustering here were three other amphipyrine genera: *Sexserrata* Barnes & Benjamin, *Tristyla* Smith, and *Metaponpneumata* Möschler. *Sexserrata* and *Tristyla* grouped sister to one another (BS=100, UF=100, SH=97.9, PP=1.00), with *Metaponpneumata* sister to *Cydosia* Duncan [& Westwood] (BS=93, UF=99, SH=97.4, PP=1.00), the lone genus in Cydosiinae. This Metoponiinae and Cydosiinae clade in turn was sister to Grotellinae (BS=88, UF=99, SH=99.5, PP=0.979). *Azenia, Tristyla, Sexserrata*, and *Metaponpneumata* are transferred to Noctuidae *incertae sedis* (see Discussion).

Four genera placed in Stiriini *incertae sedis* by Lafontaine & Schmidt (2010) were supported (BS=97, UF=100, SH=99.9, PP=1.00) as sister to the Acontiinae: *Chamaeclea* Grote, *Heminocloa* Barnes & Benjamin, *Hemioslaria* Barnes & Benjamin, and *Thurberiphaga* Dyar (Fig. 5). Chamaecleini Keegan & Wagner **New Tribe** is erected in Acontiinae for this clade of four genera which is formally described in the Discussion.

**Fig. 5.**
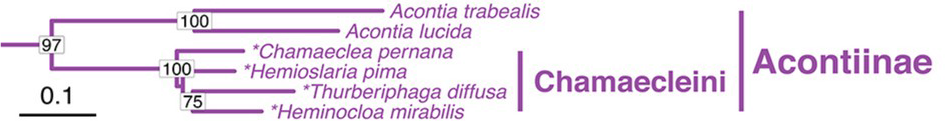
Subset of Fig 2B showing Acontiinae with tribe Chamaecleini.

Three amphipyrine genera clustered with more remote subfamilies: *Nacopa* Barnes & Benjamin was supported as sister to other Noctuinae included in this analysis (BS=100, UF=100, SH=99.6, PP=1.00), *Acopa* Harvey was supported as nesting within the Bryophilinae (BS=99, UF=100, SH=96.8, PP=1.00), and *Aleptina* Dyar was supported as sister to *Hemicephalis* Möschler (BS=100, UF=100, SH=99.4, PP=1.00) within Condicinae (Fig. 6). Male genitalic characters support these three (unexpected) results (see Discussion). *Nacopa*, *Acopa*, and *Aleptina* are transferred to Noctuinae, Bryophilinae, and Condicinae, respectively.

**Fig. 6.**
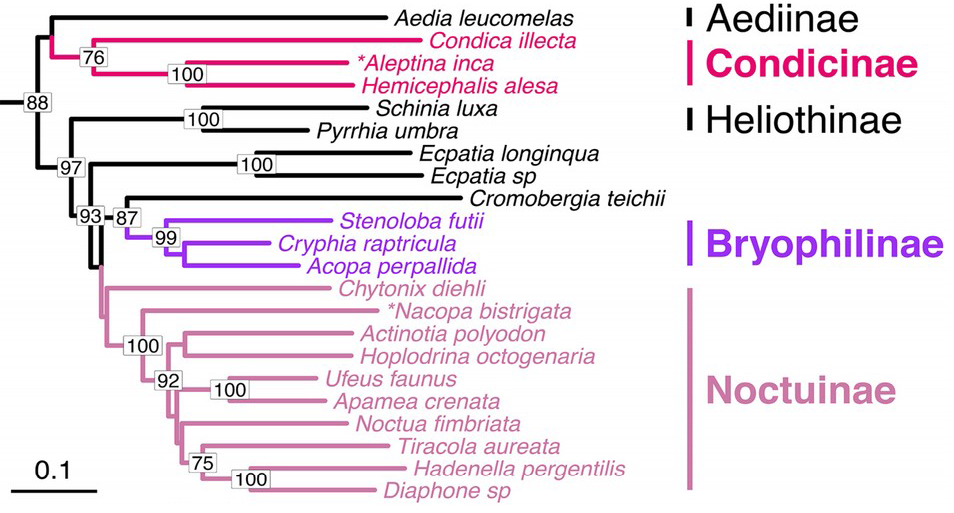
Subset of Fig 2B showing Aediinae, Condicinae, Heliothinae, Bryophilinae, and Noctuinae. The genera *Ecpatia* Turner and *Cromobergia* Bourquin are unassigned to subfamily.

A surprising finding was that *Oxycnemis* Grote contains both amphipyrines and oncocnemidines; the type species of *Oxycnemis, O. advena* Grote, clustered within Amphipyrinae s.s. (Fig. 3), whereas *O. gracillinea* (Grote) and *Leucocnemis perfundis* (Smith) clustered within Oncocnemidinae (Fig. 7) (BS=100, UF=100, SH=100, PP=1.00). *Leucocnemis* Hampson and *O. gracillinea*, but not *Oxycnemis*, are transferred to Oncocnemidinae (see Discussion).

**Fig. 7.**
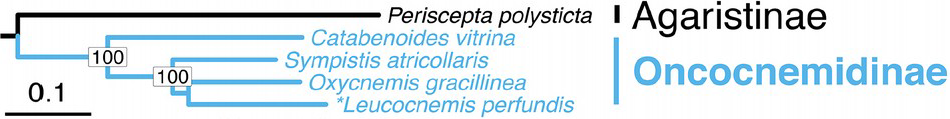
Subset of Fig 2B showing Agaristinae and Oncocnemidinae.

Also unexpected was the placement of the amphipyrine genus *Cropia* Walker which did not group with any individual subfamily. It instead grouped with the subfamilies Acronictinae through Amphipyrinae as shown in Figs 2A,B (BS=61, UF=92, SH=91, PP=0.977) with this group of subfamilies set apart as their own clade (BS=42, UF=96, SH=88.1, PP=0.97), i.e. *Cropia* was placed as the sister taxon to this massive group of taxa.

## DISCUSSION

The suspicions and misgivings of generations of workers that the Amphipyrinae were an unnatural grouping are confirmed, and staggeringly so—the 57, mostly Nearctic, amphipyrine genera surveyed fell into circa ten different subfamily-level taxa. Many taxonomic changes are needed in order to render the Amphipyrinae and other family group taxa monophyletic. Taxonomic changes (see Table 1) are recommended only for those results believed (using the seven-gene data set along with knowledge of larval morphology, adult morphology, and ecology) to be robust and unlikely to be affected by additional taxon sampling.

**Table 1.** Recommended taxonomic changes for taxa formerly regarded to be amphipyrines according to Lafontaine & Schmidt (2010, 2015).

| **Taxon** | **Current Membership** | **Recommended Change** |
| --- | --- | --- |
| *Acopa* Harvey | Amphipyridae (Psaphidini) | Bryophilinae |
| *Aleptina* Dyar | Amphipyridae (Psaphidini) | Condicinae |
| Annaphilina | Amphipyridae (Stiriini) | Stiriinae (Annaphilini) |
| *Azenia* Grote | Amphipyridae (Azeniina) | Noctuidae *incertae sedis* |
| Azeniina | Amphipyridae (Stiriini) | Noctuidae *incertae sedis* |
| *Chamaeclea* Grote | Amphipyridae (Stiriini) | Acontiinae (Chamaecleini) |
| *Cropia* Walker | Amphipyridae (Psaphidini) | Noctuidae *incertae sedis* |
| Grotellina | Amphipyridae (Stiriini) | Grotellinae |
| *Hemigrotella* Barnes & McDunnough | Amphipyridae (Stiriini) | Amphipyridae (Psaphidini) |
| *Heminocloa* Barnes & Benjamin | Amphipyridae (Stiriini) | Acontiinae (Chamaecleini) |
| *Hemioslaria* Barnes & Benjamin | Amphipyridae (Stiriini) | Acontiinae (Chamaecleini) |
| *Leucocnemis* Hampson | Amphipyridae (Psaphidini) | Oncocnemidinae |
| *Nacopa* Barnes & McDunnough | Amphipyridae (Psaphidini) | Noctuinae |
| *Nartheophora* Smith | Amphipyridae (Azeniina) | Stiriinae (Stiriini) |
| *Oxycnemis gracillinea* (Grote) | Amphipyridae (Psaphidini) | Oncocnemidinae |
| *Metaponpneumata* Möschler | Amphipyridae (Psaphidini) | Noctuidae *incertae sedis* |
| *Sexserrata* Barnes & Benjamin | Amphipyridae (Psaphidini) | Noctuidae *incertae sedis* |
| Stiriina | Amphipyridae (Stiriini) | Stiriinae |
| *Thurberiphaga* Dyar | Amphipyridae (Stiriini) | Acontiinae (Chamaecleini) |
| *Tristyla* Smith | Amphipyridae (Azeniina) | Noctuidae *incertae sedis* |

Many deeper relationships within Noctuidae (e.g. inter-subfamilial) were not well supported, as well as many subgroupings in Amphipyrinae s.s. Broader taxonomic coverage within Noctuidae and Amphipyrinae s.s., more genetic data, and/or coalescent-based phylogenetic analyses will be needed to resolve these relationships, and before formal taxonomic changes within Amphipyrinae s.s. should be made. A fuller assessment of Amphipyrinae s.s. as well as noctuid inter-subfamilial relationships is currently underway by us, with special emphasis on the subfamilies between and including Acontiinae and Amphipyrinae as shown in Fig. 2B.

Presented below are discussions of the fates of amphipyrine taxa, beginning with true Amphipyrinae (Amphipyrinae s.s.) and then moving through the amphipyrine taxa that fell outside of Amphipyrinae s.s. A limited discussion of subfamilial relationships in Noctuidae is also provided in relevant sections. For many of the tribes or subfamilies affected, a brief characterization of the morphological and life history data supporting a recommended taxonomic decision is provided.

### Amphipyrinae s.s

In large measure, the amphipyrine and psaphidine genera from Lafontaine & Schmidt’s (2010, 2015) checklist were confirmed as amphipyrines, as was the East Asian genus *Nacna*, confirming the findings of Rota *et al.* (2016). Excluded from Amphipyrinae s.s. were the entirety of Lafontaine & Schmidt’s (2010) Stiriini, which were largely dispersed among Stiriinae, Metoponiinae, and Grotellinae.

Amphipyrinae s.s. were not well supported by the RAxML analysis, but were in the other analyses. This clade was found to be well supported in previous studies based on two genes and five taxa (Mitchell *et al.*, 2006), five genes and two taxa (Regier *et al.* 2017), or eight genes and two taxa (Zahiri *et al.*, 2013). One reason for the lack of support for this group in the RAxML analysis and groupings therein might be model misspecification, as only the GTR model can be assigned to each partition in RAxML, whereas the IQ-TREE analysis explored model space and assigned the most likely model to each partition. Another potential reason for lower bootstrap support is the inclusion of multiple relatively long-branch taxa within Amphipyrinae s.s. (e.g. *Feralia* Grote, *Emarginea* Guenée, *Hemigrotella*, and *Miracavira* Franclemont), which can lower bootstrap values even for true clades (Van de Peer *et al.*, 2000).

Unlike in previous molecular studies, little support was found for the Psaphidini being monophyletic. In Europe Psaphidini are given subfamily status separate from Amphipyrinae (Fibiger & Hacker 2007). The reasons for this lack of support may well be the same as those mentioned for the lack of support of Amphipyrinae s.s.

Given the shortness of several (deeper) internal branches and weak nodal support within Amphipyrinae s.s., it would be premature to formally delimit amphipyrine tribes and subtribes before more sampling is done across amphipyrine genera (especially beyond the Nearctic Region), and/or genomic-scale data are used.

### Stiriinae

As suggested by their larvae and life histories (Crumb, 1956; Wagner *et al.*, 2011), adult morphology (Poole, 1995), and a recent molecular study of the Noctuidae (Regier *et al.*, 2017), the Stiriinae were found to represent a distinct subfamily (Figs 2B,4). As defined here, Stiriinae are trimmed relative to previous concepts (Franclemont & Todd, 1983; Poole, 1995; Lafontaine & Schmidt, 2010); restricted to what Lafontaine & Schmidt (2010) regarded as the subtribes Stiriina (with the addition of *Narthecophora*) and Annaphilina, both of which are here elevated to tribes.

Stiriinae are distributed mainly in southwestern North America, and reach greatest diversity in deserts and adjacent aridlands (Hogue, 1963). It is suspected their species and generic richness in Mexico will greatly exceed that found north of the Mexico-US border. Within Stiriini, all but a few early diverging genera are thought to be specialists on Asteraceae. Most included taxa are reliant on reproductive tissues, either flowers or callow seeds, as larvae. *Annaphila* are specialists on Boraginaceae, Montiaceae, and Phrymaceae. The subfamily is currently the focus of a species-level phylogenetic and biogeographic study by KLK.

### Grotellinae

The clade including *Grotella* Harvey, *Neogrotella* Barnes & Benjamin, and *Grotellaforma* Barnes & Benjamin (Fig. 4) is non-problematic—it is well supported by molecular, adult, larval, and life history data. Given its sister-group relationship to the clade containing Metoponiinae and Cydosiinae and relative age (branch depth), this group is recognised as a subfamily, Grotellinae, elevated from its previous rank as a subtribe. The Grotellinae are endemic to the deserts of southwestern North America and contain 23 described species (Poole, 1989). So far as known, all species are dietary specialists of Nyctaginaceae. Although several species feed on leaves, especially in early instars, most are flower and seed predators with their phenology closely tied to that of a single local host.

### Metoponiinae and Cydosiinae

This grouping of taxa (Fig. 4) is the most unorthodox and perplexing presented here. It’s unclear if the group is comprised mostly of long-branch misfits or if it is a natural, but phenotypically divergent, assemblage. Denser taxon sampling across this curious collection of genera is needed to better understand their phylogenetic relationships.

Of the seven genera treated here, the phenotypic outlier is *Cydosia*, a small, mostly tropical, genus with magnificent, highly derived larvae that seemingly set them apart from those of neighboring lineages: i.e., the prolegs on A3 and A4 are present but reduced; the D2 and SD pinacula are exceedingly elongate (sometimes > 15 × their width) on A1 and often proximate thoracic segments as well as on A2 and A3; and the apical seta on each such elongated pinaculum is lamelliform (Figs 8C,D). Early American workers commonly placed *Cydosia* in Acontiinae (McDunnough, 1938; Franclemont & Todd, 1983). Lafontaine & Schmidt (2010) transferred *Cydosia* into its own subfamily. The analyses of Zahiri *et al.* (2013) and Rota *et al.* (2016) placed *Cydosia* within Metoponiinae (rendering Metoponiinae paraphyletic in treatments that accord subfamilial rank to Cydosiinae). The results of this study reaffirm their findings and suggest four additional genera may be metoponiines: *Azenia*, *Metaponpneumata*, *Sexserrata*, and *Tristyla*.

**Fig. 8.**
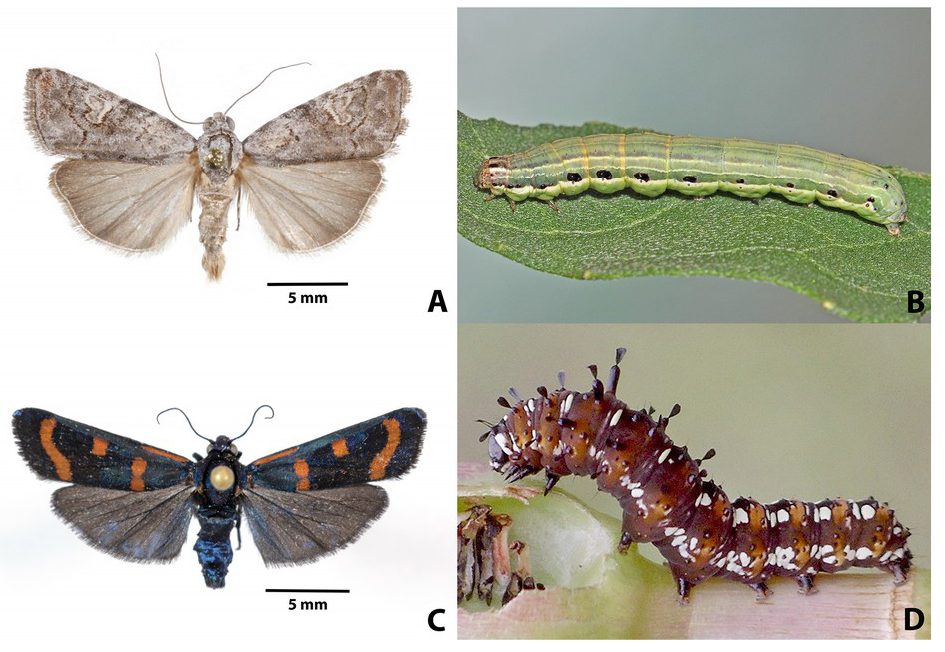
Adults and larvae of *Cydosia* and *Metaponpneumata*. A, *Metaponpneumata rogenhoferi* Möschler adult. B, *M. rogenhoferi* last instar. C, *Cydosia aurivitta* Grote & Robinson adult. D, *C. aurivitta* last instar. *(Cydosia* larval image courtesy of Valerie Bugh.)

*Metaponpneumata* and *Cydosia* grouped sister to one another. That *Cydosia* would share a recent common ancestor with *Metaponpneumata*, a small, gray, nondescript denizen of North American deserts with a similarly subdued larva (Figs 8A-D) was not expected. When *Metaponpneumata* was removed from the analysis the same topology was recovered with respect to *Flammona* Walker, *Panemeria* Hübner, and *Cydosia* (results not shown) as in Zahiri *et al.* (2013) and Rota *et al.* (2016). Interestingly, both *Cydosia* and *Metaponpneumata* are dietary generalists; so far as known other Metoponiinae are known or believed to be hostplant specialists (DLW unpublished data). Given the surprising relationships in this part of the tree, but sparse taxon sampling, no recommendations as to subfamily delineation or membership are given. Instead, these four amphipyrine genera (*Azenia, Metaponpneumata, Sexserrata, Tristyla*), along with the subtribe Azeniina, are placed into Noctuidae *incertae sedis*.

### Acontiinae and Chamaecleini

The genera *Chamaeclea*, *Heminocloa*, *Hemioslaria*, and *Thurberiphaga* formed a clade sister to the two acontiines included in the analysis: *Acontia lucida* (Hufnagel) and *Acontia trabaelis* (Scopoli) (Fig. 5). These four former amphipyrine genera are provisionally and conservatively included in the tribe Chamaecleini in Acontiinae on the basis of adult morphological characters and shared life history associations, however no characters were found in the larvae that uniquely link Chamaecleini to Acontiinae (see the description of Chamaecleini at the end of the Discussion).

### Noctuinae and Bryophilinae

Male genitalic characters support the new assignments of both *Acopa* and *Nacopa* (their larvae are unknown). The male genitalia of *Acopa* would not immediately be recognised as belonging to the Bryophilinae because the valve is short, 2 × as long as the sacculus, and heavily sclerotised, whereas in most Bryophilinae the valve is long, usually 3 × as long as the sacculus and is weakly sclerotised. Two features of the valve are similar to those found in the Bryophilinae: the uncus is flattened and slightly spatulate apically, and the clasper appears to arise from the costal margin of the valve, which is a feature common to many *Bryophila* Treitschke in Eurasia. No specimens of other New World bryophilines (e.g. “*Cryphia*” Hübner) from North America were included in this study, so the relationship of *Acopa* to the other New World representatives of the subfamily remains unclear, but some species, e.g., “*Cryphia” olivacea* (Smith), have a minute rounded clasper at the same position on the costal margin of the valve and with the same orientation as *Acopa*.

*Nacopa* has unusual valves for the subfamily Noctuinae in that the sacculus is massive, occupying about three quarters of the volume of the valve, but like other Noctuinae the clasper is high on the valve but still connected to the lower margin by the thin sclerotised band discussed by Lafontaine & Poole (1991: 21). Being placed sister to much of the rest of the Noctuinae in this study, *Nacopa* may provide evolutionary insight into early aspects of the radiation of Noctuinae—one of the most ecologically successful and economically important clades of Lepidoptera (Zhang 1994, Mitchell *et al.*, 2006).

### Condicinae

*Aleptina* was well supported as sister to *Hemicephalis* (Fig. 6). Early North American workers placed *Aleptina* in the Acontiinae (McDunnough, 1938; Franclemont & Todd, 1983; Todd *et al.*, 1984). The genus was transferred without explanation to the amphipyrine subtribe Triocnemidina by Lafontaine & Schmidt (2010). The larvae, recently revealed to be specialists on various species of *Tiquilia* (Boraginaceae) (DLW unpublished data), are consistent with other condicines, but have only two SV setae on A1, like most higher Noctuidae, and unlike other genera of Condicinae. *Aleptina* larvae resemble miniature versions of the condicine *Diastema* Guenée: the head is partially retracted into the prothorax, the prolegs on both A3 and A4 are modestly reduced, A8 is humped, and the spiracular stripe (when present) runs from the spiracle on A8 down the anal proleg. McDunnough (1938) had placed *Aleptina* and *Diastema* proximate in his checklist—a position unchanged in Franclemont & Todd (1983) and now supported by this study. Before the nuclear and mitochondrial DNA phylogeny of Zahiri *et al.* (2012) few would have thought *Hemicephalis* (previously held to be an erebid) would in fact belong in the Noctuidae, let alone the Condicinae. However, taxon sampling remains sparse in this area of the tree. Increased taxon sampling in and around this area is needed, e.g. to investigate if *Aleptina* and *Diastema* are in fact Condicinae, and not representatives of a separate known (or unknown) subfamily.

### Oncocnemidinae

Lafontaine & Schmidt (2010) placed *Leucocnemis* in the amphipyrine subtribe Triocnemidina. Its type, *Leucocnemis perfundis*, grouped with both oncocnemidines in this study (Fig. 7). Consistent with this placement, the larva has the first two pairs of prolegs greatly reduced; the setae are relatively long and borne from minute white warts; and the D2 setae on A8 arise from warts on a sharply angled, transverse ridge. The caterpillar’s fitful, prolonged alarm response is typical for oncocnemidines, but unknown from amphipyrines. Because *L. perfundis* is the type species, *Leucocnemis* is transferred to Oncocnemidinae. It is possible that some *Leucocnemis* may be triocnemidine amphipyrines.

The polyphyly found in *Oxycnemis* based on molecular data is also supported by life history data and larval characters. *Oxycnemis gracillinea*, which groups with oncocnemidines, feeds on *Menodora* (Oleaceae) (many Oncocnemidinae feed on this plant family) (Wagner *et al.*, 2011; DLW unpublished data). The caterpillar of *O. gracillinea* differs from those of *O. advena* in having no obvious rump over A8, inconspicuous dorsal pinacula, and reduced prolegs on A3 and A4—traits common to oncocnemidines. Both *O. advena* and its California cousin, *O. fusimacula* Smith, are *Krameria* (Krameriaceae) feeders. Both have a strongly humped A8, enlarged white dorsal pinacula; and full-sized anterior prolegs—traits common to amphipyrine larvae; the caterpillars also lack the thrashing alarm response of oncocnemidines. Because *O. advena* is the type species of *Oxycnemis* the genus is retained in Amphipyrinae. *O. gracillinea* is placed in Oncocnemidinae without generic assignment.

### *Cropia* Walker

*Cropia*, a Neotropical genus with 24 species (Poole 1989), fell outside of any known subfamily, and has long been recognised as an anomalous noctuid and dubious member of Amphipyrinae (Robert Poole, pers. comm.). The male genitalia of *Cropia connecta* (Smith) corroborate the molecular findings in that they are odd for Noctuidae: they are relatively large, weakly sclerotised, and set with a curious abundance of soft piliform setae. *C. connecta*, the sole representative of the genus in this study, has genitalia substantially different from those of the type species, *C. hadenoides* Walker. The larva of *C. hadenoides* also differs markedly from other species in the genus (Dan Janzen pers. comm.). Given the possibility that *Cropia* may represent two distinct lineages, no subfamily assignment of *Cropia* is recommended other than its removal from Amphipyrinae and placement in Noctuidae *incertae sedis*.

### Taxonomy

#### Chamaecleini Keegan & Wagner, 2018 New Tribe (Noctuidae, Acontiinae)

http://zoobank.org/urn:lsid:zoobank.org:act:0D86A34B-AB52-4114-B6C7-1EC2953D0175

#### Type genus

*Chamaeclea* Grote, 1883.

#### Type species

*Chariclea pernana* Grote, 1881.

#### Diagnosis

Chamaecleini differ from other tribes of the Acontiinae in having scattered setae on the scaphium, not clustered into a tuft or tufts of setae; claspers symmetrical or very slightly asymmetrical, not markedly asymmetrical; larvae with prolegs on A3-A6 and without modified anal setae of Acontiini.

#### Adult Description

Characters in **bold** distinguish Chamaecleini from other Acontiinae, characters in *italics* are shared with and apomorphic for Acontiinae.

<u>Head</u>: antenna of male and female filiform, scaled dorsally; laterally and ventrally unscaled and densely pubescent with minute setae; **frons with frontal tubercle consisting of raised rounded ring, open ventrally, with low conical tubercle in center**; eye rounded, smooth; palpi porrect, scaled, without tufts; haustellum functional, coiled. <u>Thorax</u>: prothoracic collar and thorax clothed with spatulate, apically serrated, scales; forewing with typical noctuid quadrifine venation (i.e., vein M_2_ close to M_3_); hind wing venation trifine (M_2_ reduced, slightly closer to M_3_ than to M_1_; legs typical of most Noctuidae (without spine-like setae on tibiae, and without spine at apex of foretibia); *tympanal opening with hood vestigial*, *and alula enlarged and clothed with large flat scales that cover 1/3-2/3 of opening*; **tympanal sclerite a sclerotised ridge with surface only slightly nodular**, unlike nodular sclerite of most higher Noctuidae. <u>Abdomen</u>: long slender apodemes on basal sternite; without basal hair-pencils, levers, or pockets. <u>Male genitalia</u>: uncus slender, sparsely setose, curved downward to pointed apex; tegumen broad, tapered abruptly ventrad, connected to vinculum by broad plural sclerite fused to vinculum; vinculum broadened ventrally into U-shaped saccus; scaphium mainly membranous, lightly sclerotised ventrally, **with scattered short setae dorsally**, **not clustered into one or two patches as in Acontiini**; **valves symmetrical**; sacculus extending from valve base 1/3 of distance to valve apex and differentiated from clasper only by lightly sclerotised junction; clasper broad basally with more heavily-sclerotised lobe on dorsum and ending in small rounded lobe near valve apex; valve with no apical corona of heavily sclerotised setae; aedeagus 5 × as long as wide; vesica slightly longer than aedeagus with ventral and subbasal pouches with spinules on subbasal pouch and near vesica apex. <u>Female genitalia</u>: Anal papillae long and tapered to apex, clothed with short setae; posterior and anterior apophyses long 4 × and 3 × as long as abdominal segment 8; ductus bursae 3 × as long as abdominal segment 8, lightly sclerotised posteriorly; corpus bursae very long and slightly coiled, 16 × as long as abdominal segment 8. Tapered anal papillae and elongated apophyses suggest telescoping oviposition, probably into flowers.

#### Larval Description

Characters in **bold** distinguish Chamaecleini from other Acontiinae. **Fully legged with well-developed**, **crochet-bearing prolegs on A3-A6**; **dorsal and ventral anal comb setae described by Crumb (1956) lacking; spinneret elongate; SV1 is well forward of SV2 and SV3**. Feed on seeds and flowers of Malvaceae.

#### Included Taxa

*Chamaeclea* includes two species with *C. basiochrea* Barnes & McDunnough from Texas being similar both in external appearance and in genital characters to *C. pernana*. In addition to *Chamaeclea*, the Chamaecleini include four monobasic genera that differ from *Chamaeclea* in the following: ***Heminocloa mirabilis*** (Neumoegen) [setae on scaphium long, hair-like; male valve strap-like; clasper heavily sclerotised and well differentiated from sacculus, with dorsal process in middle and pointed apical process free from valve]; female genitalia not examined. ***Hemioslaria pima*** Barnes & Benjamin [only a few minute setae on scaphium; male genitalia similar to those of *H. mirabilis*, except valve almost triangular due to large dorsal lobe; clasper without dorsal process; vesica globular]; female genitalia not examined. ***Thurberiphaga diffusa*** (Barnes) [antenna lamellate, branches longer in males than females; clasper fused into valve, made evident mainly by series of setae on bumps along middle of valve; female genitalia with anal papillae short, pad-like and densely setose; apophyses and ductus bursae relatively short; corpus bursae about 4 × as long as abdominal segment 8 and pear shaped]. Larvae are known for three genera: *Chamaeclea*, *Heminocloa*, and *Thurberiphaga*. All feed on Malvaceae as do most Acontiinae (Crumb, 1956; Wagner *et al.*, 2011; DLW unpublished data). The smooth, grub-like caterpillars bore into ripening fruits to feed on seeds—a far less common feeding strategy than leaf feeding among acontiines (Crumb, 1956; DLW unpublished data). Larval characters for *Heminocloa* and *Thurberiphaga*, as given above, except spinneret long-enough to bear lateral sclerites in both genera.

#### Remarks

Although molecular, adult genital and tympanal characters, and life history data suggest a sister group relationship between Acontiini and Chamaecleini, no larval characters were found that were uniquely shared with Acontiini, i.e. none of the characters in Crumb’s (1956) larval key to noctuid subfamilies apomorphic for Acontiinae is expressed in the known larvae of Chamaecleini. Crumb (1956) treated the larva of *Thurberiphaga*, but left it unassigned to any subfamily.

## CONCLUSION

The realm of Amphipyrinae has waxed and waned for more than a century, with no two major taxonomic works seeming to agree on the limits of the subfamily. More expansive concepts have spanned the subfamilies that were the focus of this study (e.g., Edwards, 1996) whereas others restricted its content to just the nominate genus (e.g., Kitching & Rawlins, 1998). In most checklists and faunal works Amphipyrinae served as a repository for noctuids that lacked the synapomorphies of acontiines, acronictines, bagisarines, eustrotiines, cuculliines, oncocnemidines, plusiines, and others. This contribution is a step forward and provides phylogenetic scaffolding around which future taxonomic and phylogenetic efforts can be built.

Future efforts are needed to add more Old World taxa, especially from East Asia and the southern Hemisphere, and much remains to be done with the fauna of North America. Central and northern Mexico could prove to be the cradle for much the New World diversity of Amphipyrinae, Grotellinae, Metoponiinae, and Stiriinae. The type species for more than a dozen genera included in the Amphipyrinae by Lafontaine & Schmidt (2010) have yet to be sampled, and it is not improbable that other amphipyrines, unrecognised as such, still reside within other subfamilies. In addition to Amphipyrinae, the monophyly of other subfamilies (e.g., Metoponiinae, Oncocnemidinae, and Stiriinae) were also revealed to be in need of closer scrutiny. Some taxa (e.g. Cydosiinae) were shown to potentially be poor candidates for subfamilial rank, whereas others were found to be perhaps worthy of subfamilial status (e.g. *Cropia* and Chamaecleini).

As noted above, the seven genes used resolved relationships within virtually every subfamily-level taxon, but frustratingly only modest or ambiguous support for the phylogenetic relationships among the various noctuid subfamilies—a finding that supports the suggestions of others that the early radiation of the Noctuidae was a rapid one (Wahlberg *et al.*, 2013, Zahiri *et al.*, 2013). Adding more taxa and/or more genes may help clarify inter- and intra-subfamilial relationships in Noctuidae; likewise coalescent-based phylogenetic inference methods should help combat the confounding effects of incomplete lineage sorting that tend to plague rapid radiations.

As much as possible type species were emphasised in this assessment because it was evident at the outset that several amphipyrine s.l. genera were polyphyletic, such as *Oxycnemis* and *Leucocnemis*. Other genera that appear to be unnatural assemblages include *Aleptina*, *Azenia*, *Nocloa* Smith, *Paratrachea* Hampson, *Paramiana* Barnes & Benjamin, and *Plagiomimicus* Grote.

It is hoped that the relationships hypothesised in this work will facilitate efforts to identify further morphological and life history data that can be used to corroborate or refute the relationships presented in Figs 2A,B. Given the weak support for some clades, larval, anatomical, behavioral, and life history details could do much to test this study’s findings. Even in those cases where support is strong, such information is needed to add biological meaning to the inferred clades and their taxonomic concepts.

## ACKNOWLEDGEMENTS

Many of the taxa in this study are desert dwellers, tied to the stochasticity of desert rains, and as such are immensely difficult for one person to collect in a single year or even lifetime. The authors owe an immense debt to those who provided amphipyrine s.l. taxa. Three were especially generous with their specimens, time, and unpublished data: John Palting, Evan Rand, and David Wikle. The latter made a special effort to fill taxonomic gaps in preliminary analyses. Additional specimens used here were supplied by Robert Behrstock, John DeBenedictis, Cliff Ferris, Ann Hendrickson, Sam Jaffe, Ed Knudson, and Eric Metzler. Sequence data were interwoven into a pre-existing latticework built by Reza Zahiri and co-workers. Marek Borowiec assisted with initial sequence alignment and an initial phylogenetic analysis. Kennedy Marshall helped with artwork and figure design. Jim Dice, Gina Moran, and the California Department of Food and Agriculture assisted with collecting permits for Anza-Borrego Desert State Park in 2016 and 2017; the Steele-Burnand Desert Research Institute hosted KLK, DLW, and others in both years. The authors also thank Jonena Hearst and Raymond Simpson for assisting with permitting and logistics for Big Bend and Guadalupe Mountains National Parks. Steven Passoa (USDA/APHIS/ PPQ) assisted in morphological analysis of Chamaecleini larvae. The authors are grateful to Charles Mitter and two other referees for helpful comments on the manuscript. KLK is supported by grants from the Society of Systematic Biologists (Graduate Student Research Award), the Department of Ecology and Evolutionary Biology at the University of Connecticut (Zoology Award), as well as many generous donors that contributed to an Instrumentl [sic] crowd-funding grant. NW acknowledges support from the Swedish Research Council. DLW is supported by USFS Co-op Agreement 14-CA-11420004-138 and an award from the Richard P. Garmany Fund (Hartford Foundation for Public Giving).

## CONFLICT OF INTEREST

The authors declare no conflict of interest.

## REFERENCES

Altschul, S.F., Gish, W., Miller, W., Myers, E.W. & Lipman, D.J. (1990) Basic local alignment search tool. Journal of Molecular Biology, 215, 403–410.

Borowiec, M.L. (2016) AMAS: a fast tool for alignment manipulation and computing of summary statistics. PeerJ, 4, e1660.

Chernomor O., von Haeseler A. & Minh B.Q. (2016) Terrace aware data structure for phylogenomic inference from supermatrices. Systematic Biology, 65, 997–1008.

Cho, S., Mitchell, A., Regier, J.C., Mitter, C., Poole, R.W., Friedlander, T.P. & Zhao, S. (1995) A highly conserved nuclear gene for low-level phylogenetics: elongation factor-1a recovers morphology-based tree for heliothine moths. Molecular Biology and Evolution, 12, 650–656.

Crumb, S.E. (1956) The Larvae of the Phalaenidae. Technical Bulletin # 1135, United States Department of Agriculture, Washington, D.C., USA.

Edwards, E.D. (1996) Noctuidae. Checklist of the Lepidoptera of Australia (ed. by E.S. Nielsen, E. D. Edwards and T. V. Rangsi), pp. 291–333. CSIRO Publishing, Collingwood, Victoria, Australia.

Fang, Q.Q., Cho, S., Regier, J.C., Mitter, C., Matthews, M., Poole, R.W., Friedlander, T.P. & Zhao, S. (1997) A new nuclear gene for insect phylogenetics: dopa decarboxylase is informative of relationships within Heliothinae (Lepidoptera: Noctuidae). Systematic Biology, 46, 269–283.

Fibiger, M. & Hacker, H. (2004) Systematic list of the Noctuoidea of Europe. Esperiana, 11, 83–172.

Fibiger, M. & Hacker, H. (2007) Noctuidae Europaeae, Amphipyrinae, Condicinae, Eriopinae, Xyleninae (Part), Vol. 9. Entomological Press, Sorø, Denmark.

Fibiger, M. & Lafontaine, J.D. (2005) A review of the higher classification of the Noctuoidea (Lepidoptera), with special reference to the Holarctic fauna. Esperiana, 11, 7–92.

Franclemont, J.G. & Todd, E.L. (1983) Noctuidae. Check List of the Lepidoptera of America North of Mexico, (ed. by R.W. Hodges et al), pp. 120–159. E.W. Classey Ltd. and The Wedge Entomological Research Foundation, Cambridge University Press, Cambridge, UK.

Guindon, S., Dufayard, J.-F., Lefort, V., Anisimova, M., Hordijk, W. & Gascuel, O. (2010) New algorithms and methods to estimate maximum-likelihood phylogenies: assessing the performance of PhyML 3.0. Systematic Biology, 59, 307–321.

Hampson, G.F. (1898-1913) Catalogue of the Lepidoptera Phalaenae in the British Museum, British Museum Order of the Trustees, London, UK.

Hillis, D.M. & Bull, J.J. (1993) An empirical test of bootstrapping as a method for assessing confidence in phylogenetic analysis. Systematic Biology, 42,182–192.

Hoang, D.T., Chernomor, O., von Haeseler, A., Minh, B.Q. & Vinh, L.S. (2018) UFBoot2: improving the ultrafast bootstrap approximation. Molecular Biology and Evolution, 35, 518–522.

Hogue, C.L. (1963) A definition and classification of the tribe Stiriini (Lepidoptera: Noctuidae). Los Angeles County Museum Contributions in Science, 64, 1–129.

Kalyaanamoorthy, S., Minh, B.Q., Wong, T.K.F., von Haeseler, A. & Jermiin, L.S. (2017) ModelFinder: fast model selection for accurate phylogenetic estimates. Nature Methods, 14, 587–589.

Kearse, M., Moir, R., Wilson, A., Stones-Havas, S., Cheung, M., Sturrock, S., Buxton, S., Cooper, A., Markowitz, S., Duran, C., Thierer, T., Ashton, B., Mentjies, P. & Drummond, A. (2012) Geneious basic: an integrated and extendable desktop software platform for the organization and analysis of sequence data. Bioinformatics, 28, 1647–1649.

Kitching, I.J. (1984) An historical review of the higher classification of the Noctuidae (Lepidoptera). Bulletin of the British Museum (Natural History) (Entomology), 49, 153–234.

Kitching, I.J. & Rawlins, J.E. (1998) The Noctuoidea. Lepidoptera, Moths and Butterflies. Vol.1: Evolution, Systematics, and Biogeography, Handbook for Zoology, Vol. IV: Arthropoda: Insecta (ed. N.P. Kristensen), pp. 355–401. Walter de Gruyter, Berlin, Germany.

Lafontaine, J.D. & Poole, R.W. (1991) Noctuoidea, Noctuidae (Part) - Plusiinae. The Moths America north of Mexico, Fascicle 25.1 (ed. by R.W. Hodges et al.). The Wedge Entomological Research Foundation, Washington, D.C., USA.

Lafontaine, J.D. & Schmidt, B.C. (2010) Annotated check list of the Noctuoidea (Insecta, Lepidoptera) of North America north of Mexico. ZooKeys, 40, 1–239.

Lafontaine, J.D. & Schmidt, B.C. (2015) Additions and corrections to the check list of the Noctuoidea (Insecta, Lepidoptera) of North America north of Mexico III. ZooKeys, 527, 127–147.

Larsson, A. (2014) AliView: a fast and lightweight alignment viewer and editor for large datasets. Bioinformatics, 30, 3276–3278.

McDunnough, J.H. (1938) Checklist of the Lepidoptera of Canada and the United States of America. Part 1. Macrolepidoptera. Memoirs of the Southern California Academy of Sciences, 1, 1–275.

Miller, M.A., Pfeiffer, W. & Schwartz, T. (2010) Creating the CIPRES Science Gateway for inference of large phylogenetic trees. Proceedings of the Gateway Computing Environments Workshop (GCE), New Orleans, Louisiana, 14 November 2010.

Mitchell, A., Mitter, C. & Regier, J.C. (2006) Systematics and evolution of the cutworm moths (Lepidoptera: Noctuidae): evidence from two protein-coding nuclear genes: molecular systematics of Noctuidae. Systematic Entomology, 31, 21–46.

NCBI Resource Coordinators. (2017) Database resources of the National Center for Biotechnology Information. Nucleic Acids Research, 45, D12–D17.

Nguyen, L.-T., Schmidt, H.A., von Haeseler, A. & Minh, B.Q. (2015) IQ-TREE: a fast and effective stochastic algorithm for estimating maximum-likelihood phylogenies. Molecular Biology and Evolution, 32, 268–274.

Poole, R.W. (1989) Lepidopterorum Catalogus (New Series). Fascicle 118. Noctuidae [in 3 Parts] E.J. Brill/Flora and Fauna Publications, New York, USA.

Poole, R.W. (1995) Noctuoidea. Noctuidae (part). Cuculliinae, Stiriinae, Psaphidinae (Part). The Moths of America North of Mexico, Fascicle 26.1 (ed. by R. B. Dominick et al.)., Washington, D.C., USA.

R Core Team. (2017) R: A language and environment for statistical computing. R Foundation for Statistical Computing, Vienna, Austria. https://www.R-project.org/.

R Studio Team. (2015) RStudio: Integrated Development for R. RStudio, Inc., Boston, MA http://www.rstudio.com/.

Rambaut, A., Suchard, M.A., Xie D. & Drummond, A.J. (2014) Tracer v1.6, Available from http://tree.bio.ed.ac.uk/software/tracer/.

Ratnasingham, S. & Hebert, P.D.N. (2007) BOLD: the barcode of life data system (www.barcodinglife.org). Molecular Ecology Notes, 7, 355–364.

Regier, J.C., Mitter, C., Mitter, K., Cummings, M.P., Bazinet, A.L., Hallwachs, W., Janzen D.H. & Zwick, A. (2017) Further progress on the phylogeny of Noctuoidea (Insecta: Lepidoptera) using an expanded gene sample. Systematic Entomology, 42, 82–93.

Ronquist, F., Teslenko, M., van der Mark, P. et al. (2012) MrBayes 3.2: efficient Bayesian phylogenetic inference and model choice across a large model space. Systematic Biology, 61, 539–542.

Rota, J., Zacharczenko, B.V, Wahlberg, N., Zahiri, R., Schmidt, B.C. & Wagner, D.L. (2016) Phylogenetic relationships of Acronictinae with discussion of the abdominal courtship brush in Noctuidae (Lepidoptera). Systematic Entomology, 41, 416–429.

Stamatakis, A., Hoover, P., Rougemont, J. & Renner, S. (2008) A rapid bootstrap algorithm for the RAxML web servers. Systematic Biology, 57, 758–771.

Todd, E.L., Blanchard, A. & Poole, R.W. (1984) A revision of the genus Aleptina (Lepidoptera: Noctuidae). Proceedings of the Entomological Society of Washington, 86, 951–960.

Trifinopoulos, J. & Minh, B. (2018) IQ-TREE Manual: Frequently Asked Questions. Available from http://www.iqtree.org/doc/Frequently-Asked-Questions. Accessed 18 July 2018.

Troubridge, J.T. (2008) A generic realignment of the Oncocnemidini sensu Hodges (1983) (Lepidoptera: Noctuidae: Oncocnemidinae), with descriptions of a new genus and 50 new species. Zootaxa, 1903, 1–95.

Van de Peer, Y., Baldaufrid, S.L., Doolittle, W.F. & Meyerid, A. (2000) An updated and comprehensive rRNA phylogeny of (crown) eukaryotes based on rate-calibrated evolutionary distances. Journal of Molecular Evolution, 51, 565–576.

Wagner, D.L., McFarland, N., Lafontaine, J.D. & Connolly, B. (2008) Early stages of Miracavira brillians (Barnes) and reassignment of the genus to the Amphipyrinae: Psaphidini: Feraliiina (Noctuidae). Journal of the Lepidopterists’ Society, 62, 121–132.

Wagner, D.L., Schweitzer, D.F., Sullivan, J.B. & Reardon, R.C. (2011) Owlet Caterpillars of Eastern North America. Princeton University Press, Princeton, New Jersey, USA.

Wahlberg, N. & Wheat, C.W. (2008) Genomic outposts serve the phylogenomic pioneers: designing novel nuclear markers for genomic DNA extractions of Lepidoptera. Systematic Biology, 57, 231–242.

Wahlberg, N., Wheat, C.W. & Peña, C. (2013) Timing and patterns in the taxonomic diversification of Lepidoptera (butterflies and moths). PLoS ONE, 8, e80875.

Yu, G., Smith, D.K., Zhu, H., Guan, Y. & Lam, T.T.Y. (2017) ggtree: an R package for visualization and annotation of phylogenetic trees with their covariates and other associated data. Methods in Ecology and Evolution, 8, 28–36.

Zahiri, R., Kitching, I.J., Lafontaine, J.D., Mutanen, M., Kaila, L., Holloway, J.D. & Wahlberg, N. (2011) A new molecular phylogeny offers hope for a stable family level classification of the Noctuoidea (Lepidoptera). Zoologica Scripta, 40, 158–173.

Zahiri, R., Holloway, J.D., Kitching, I.J., Lafontaine, J.D., Mutanen, M. & Wahlberg, N. (2012) Molecular phylogenetics of Erebidae (Lepidoptera, Noctuoidea). Systematic Entomology, 37, 102–124.

Zahiri, R., Lafontaine, D., Schmidt, C., Holloway, J.D., Kitching, I.J., Mutanen, M. & Wahlberg, N. (2013) Relationships among the basal lineages of Noctuidae (Lepidoptera, Noctuoidea) based on eight gene regions. Zoologica Scripta, 42, 488–507.

Zhang, B.C. (1994) Index of Economically Important Lepidoptera. CAB International, Wallingford, UK.

